# Colocalization and discordance between plasma and brain protein quantitative trait loci

**DOI:** 10.64898/2026.05.01.722237

**Authors:** Yanzhen Cheng, Wenmin Zhang, Tianyuan Lu

## Abstract

Studies of protein quantitative trait loci (pQTLs) provide opportunities to interpret complex trait genetics and identify potential biomarkers and therapeutic targets. Circulating proteins are commonly used in pQTL studies due to the accessibility of blood-based measurements, but their levels may not always reflect regulation in disease-relevant tissues. We assessed colocalization and discordance between plasma and dorsal prefrontal cortex cis-pQTLs using data from four large-scale studies and investigated their implications for downstream analyses. Across the proteins examined, at most 80% of the cis-pQTLs showed evidence of colocalization. Among the colocalized loci, approximately 20% exhibited opposite directions of genetic effects. We characterized tissue-specific gene expression profiles based on data from the Genotype-Tissue Expression project. Proteins with colocalized cis-pQTLs were more likely to have high gene expression levels in systemic tissues and immune cells, whereas the remaining proteins were more likely to have high expression in brain tissues. We conducted Mendelian randomization (MR) analyses using neuroticism as an illustrative outcome to compare effect estimates derived using instruments from different pQTL studies. MR analyses identified 13 proteins significantly associated with neuroticism, including six with opposite effect directions between plasma and dorsal prefrontal cortex, highlighting the importance of tissue context. Overall, circulating pQTLs remain informative for proteins from systemic and immune pathways, while incorporating tissue-specific data may provide additional insight for proteins with more localized expression. Considering multiple tissue contexts may refine the interpretation of protein-trait associations and may improve the prioritization of candidate targets.

## Introduction

Proteins play central roles in nearly all biological processes, acting as enzymes, signaling molecules, structural components, and regulators of cellular function^1–3^. Understanding the mechanisms that regulate protein abundance is therefore critical for elucidating disease biology and identifying potential therapeutic targets. In recent years, studies of protein quantitative trait loci (pQTLs), which identify genetic variants associated with protein abundance, have provided a powerful framework for understanding the genetic regulation of proteins^4^. By integrating pQTL data with genetic association studies of complex traits, approaches such as Mendelian randomization (MR) have enabled the systematic identification of candidate biomarkers and potential drug targets^4–8^.

Circulating proteins have attracted particular attention in this context. Blood-based proteomic measurements are relatively accessible and scalable, making them well suited for large population studies. Several large-scale efforts, including the deCODE study, the Fenland study, and the UK Biobank Pharma Proteomics Project (UKB-PPP), have generated comprehensive maps of circulating pQTLs in tens of thousands of individuals ^9–11^. These resources have enabled large-scale genetic analyses of the plasma proteome and have facilitated numerous MR studies evaluating the potential roles of circulating proteins in complex diseases.

Despite these advances, circulating protein abundance may not accurately reflect biologically active protein levels in relevant tissues^12^. Many proteins measured in plasma originate from multiple tissues and may enter the circulation through secretion, cellular turnover, or tissue injury^13,14^. Consequently, genetic variants associated with circulating protein concentrations may not capture the regulation of proteins within the tissues where they exert their primary biological functions. The direction or magnitude of genetic effects on circulating protein levels may differ from those in relevant tissues^15,16^. These considerations highlight the importance of understanding the extent to which circulating pQTLs reflect the underlying genetic regulation of proteins in disease-relevant tissues.

Directly addressing this issue has been challenging because large-scale proteomic studies of human tissues remain limited. Compared with plasma proteomics, tissue-specific proteomic profiling is considerably more expensive and technically demanding^17^, and large cohorts with both proteomic and genetic data are rare^18^. Recently, a large-scale pQTL study of the dorsolateral prefrontal cortex generated proteomic and genetic data from over 1,000 post-mortem brain samples^19^. Although proteins detected in these samples may also originate from multiple sources, including local cellular production and proteins transported from other tissues, this dataset provides a unique opportunity to systematically evaluate the extent to which circulating pQTL signals overlap with those observed in brain tissues.

In this study, we leveraged this brain pQTL resource together with three large circulating pQTL datasets from the deCODE study, the Fenland study, and the UKB-PPP to investigate the genetic regulation of protein abundance in plasma and brain. First, systematic colocalization analyses were conducted to assess the extent to which pQTL signals are shared between circulating and brain proteomes. Second, using gene expression data from the Genotype-Tissue Expression (GTEx) project^20^, tissue expression patterns were examined for proteins whose pQTL signals did and did not colocalize across tissues, providing insights into the tissues where these proteins may be produced. Finally, MR analyses were performed with neuroticism as an illustrative outcome. Genetic instruments that colocalized between plasma and brain pQTLs were used to evaluate whether consistent estimates of potential protein effects on neuroticism could be obtained across tissues. By characterizing the degree of overlap and potential discrepancies between circulating and brain pQTL signals, this work highlights the importance of considering tissue context when interpreting genetic associations with circulating proteins and evaluating their potential roles in complex traits.

## Method

### Dorsolateral prefrontal cortex protein measurements

Proteomic and genetic data from dorsolateral prefrontal cortex tissue were obtained from large collaborative efforts within the Accelerating Medicines Partnership-Alzheimer’s Disease (AMP-AD)^21^ and AMP-AD Diversity programs^22,23^. Brain samples were collected by multiple research centers, including the Rush Alzheimer’s Disease Center, Mayo Clinic, Mount Sinai Hospital, Emory University, and the Brain and Body Donation Program at Banner Sun Health^19^. All donors or their next of kin provided informed consent, and study protocols were approved by the institutional review boards at the participating institutions. After quality control procedures for both genetic and proteomic data, a total of 1,013 non-Hispanic White individuals remained for pQTL identification^19^.

### Circulating protein measurements

Circulating pQTL data were obtained from three large population studies consisting of participants of European ancestry: the deCODE study (n = 35,559)^11^, the Fenland study (n = 10,708)^10^, and the UKB-PPP (n = 34,557)^9^.

Plasma protein concentrations in the deCODE and Fenland studies were measured using the SomaScan v4 aptamer-based proteomic platform. Standard SomaLogic normalization procedures were applied, including hybridization normalization, median signal normalization across samples, and calibration across assay plates. Samples or protein measurements that failed platform quality control were excluded prior to downstream analyses^10,11^. In the UKB-PPP dataset, proteomic measurements were generated using the Olink proximity extension assay platform. Raw counts were converted to normalized protein expression values using Olink’s MyData Cloud software. Built-in internal controls, including incubation, extension, and amplification controls, were used to monitor assay performance and correct for technical variation. Additional sample-level and plate-level quality control procedures were applied to ensure reliable measurements^9^.

Details of these studies, including pQTL identification procedures, have been described previously^9–11^. For all datasets, we focused on cis-pQTLs, as these variants are more likely to have direct regulatory effects on protein expression and are less prone to pleiotropic or distal confounding compared to trans-pQTLs. In addition, this focus ensured consistency across datasets, as only cis-pQTL summary statistics were available for the brain pQTL study. Genetic variants located within 500 kb of the protein-coding gene were considered cis variants and were used for downstream analyses.

### Colocalization analysis

To evaluate whether pQTL signals observed in circulating proteomes and brain tissue reflect the same underlying causal variants, we performed colocalization analyses using SharePro^24^. SharePro estimates the probability that two traits share a causal variant while accounting for linkage disequilibrium (LD) and the presence of one or multiple causal signals, and has demonstrated improved power to identify shared causal signals while maintaining appropriate false positive control. Specifically, SharePro uses a sparse projection framework to group correlated variants into credible sets representing independent association signals. An efficient variational inference algorithm then jointly infers both the variant representations and causal status of each credible set. Colocalization probabilities are then obtained from the posterior probabilities that the same credible set is causal for both traits^24^.

For each pair of plasma and brain pQTLs, we retained cis variants that were present in both pQTL summary statistics and had a minor allele frequency > 0.05. We retained proteins that had genome-wide significant genetic associations (p-value < 5×10^-8^) in both the brain pQTL study and at least one circulating pQTL study. Otherwise, a lack of colocalization might reflect limited statistical power rather than true differences in genetic associations^25^. We used 5,000 randomly selected UK Biobank individuals of European ancestry to construct an LD reference panel^26^. Analyses were performed using the default prior settings of SharePro, and maximum colocalization probabilities were evaluated across configurations allowing one (K = 1) or up to ten (K = 10) credible sets^24^. A colocalization probability greater than 0.8 was considered evidence of colocalization^5–8,24^. For each pair of colocalized pQTLs, we further examined whether the effect direction was consistent between plasma and brain by comparing the pQTL effect estimates of the lead cis-pQTL variant with the highest posterior inclusion probability.

### Quantification of tissue-specific gene expression

To characterize tissue expression patterns of proteins with and without colocalized pQTL signals, we analyzed gene expression data from the GTEx project (version 8)^20^, which includes RNA-sequencing data across 54 non-diseased human tissues. We applied an established approach to normalize gene expression levels across samples and tissues^27^.

Genes were first filtered to retain those detected in at least 20% of samples with a minimum of five read counts. Expression values were normalized within each tissue using trimmed mean of M-values normalization implemented in the edgeR R package (version 4)^28^. For each gene and tissue, we calculated the median normalized expression across individuals.

Within each tissue, the distribution of median expression values across genes was standardized using the median and the median absolute deviation. Subsequently, tissue-specific enrichment scores for each gene were obtained by standardizing these values across tissues using the same statistics. Tissues with enrichment z-scores above 2 were considered to show elevated tissue-specific expression for the corresponding gene.

For each GTEx tissue, we performed Chi-squared tests to examine whether proteins with colocalized circulating and dorsolateral prefrontal cortex pQTL (colocalization probability > 0.8) were more or less likely to show elevated tissue-specific expression.

### Mendelian randomization analysis

To evaluate whether discrepancies between plasma and brain pQTL impact downstream genetics-guided biomarker and drug target discovery efforts, we performed MR analyses using neuroticism as the outcome. We focused on neuroticism because the dorsolateral prefrontal cortex is crucial for higher-order cognitive processing and emotional regulation^29^, which are closely related to personality traits.

Genome-wide association study (GWAS) summary statistics for neuroticism were obtained from a meta-analysis of neuroticism scores measured in the Million Veteran Program, UK Biobank, 23andMe, and Genetics of Personality Consortium comprising up to 682,688 individuals of predominantly European ancestry^30,31^.

Proteins with colocalized plasma and brain pQTL were included as exposures. We did not focus on proteins without colocalized pQTL or proteins for which pQTL was identified only in plasma or brain, because for those proteins, differences in MR estimates could reflect limited statistical power in the pQTL studies rather than true differences in genetic associations. For each colocalized credible set, we selected the lead cis-pQTL variant with the highest posterior inclusion probability inferred by SharePro to serve as the genetic instrument, ensuring independence between instruments.

For proteins with a single genetic instrument, MR estimates were calculated using the Wald ratio^32^. For proteins with two instruments, inverse-variance weighted estimates^33^ were calculated. Because no significant association was detected based on three or more independent instruments, additional sensitivity analyses using alternative MR methods were not performed.

Two-sample MR analyses were conducted using the TwoSampleMR R package (version 0.6.30)^34^.

Statistical significance was defined as a p-value < 1×10^-5^. This threshold corresponds to a Bonferroni-corrected threshold considering 5,000 proteins and may be conservative because of correlations among proteins and potential functional relationships. However, this stringent threshold provides effective control of false positives.

## Results

### Colocalization of dorsolateral prefrontal cortex pQTLs and circulating pQTLs

Table 1 summarizes the four proteomic cohorts included in this study. Notably, dorsolateral prefrontal cortex proteomic data were generated using mass spectrometry and underwent normalization procedures that differ from those applied to circulating proteomic data. Consequently, the magnitudes of genetic associations identified in the brain pQTL study may not be directly comparable to those in the circulating pQTL studies, and downstream analyses therefore focused on comparing effect directions.

**Table 1.** Summary of pQTL studies.

| Source | Dorsolateral prefrontal cortex pQTL<sup>19</sup> | Circulating pQTL in deCODE<sup>11</sup> | Circulating pQTL in Fenland<sup>10,35</sup> | Circulating pQTL in UKB-PPP<sup>9</sup> |
| --- | --- | --- | --- | --- |
| Sample size | n = 1,013 post-mortem samples collected from | n = 35,559 European ancestry | n = 10,708 European ancestry individuals | n = 34,557 European ancestry individuals |
|  | non-Hispanic White individuals in United States | individuals from Iceland | from United Kingdom | from United Kingdom |
| Platform | Tandem mass tag mass spectrometry | Aptamer-based SomaScan assay v4 | Aptamer-based SomaScan assay v4 | Antibody-based Olink Explore 3072 proximity extension assay |
| Number of proteins measured | 9,725 | 4,907 | 4,979 | 2,923 |
| GWAS procedures | Genome-wide association analyses were performed on normalized protein abundance. Protein levels were first normalized for total abundance and log<sub>2</sub>-transformed, followed by regression on covariates and surrogate variable analysis to account for unobserved confounding. Residuals were then standardized to Z-scores. Analyses were adjusted for sex, and relatedness and population structure were accounted for using a linear mixed model implemented in GEMMA. | Protein levels were first adjusted for age, sex, and sample age. The residuals were then rank-inverse normal transformed, and these transformed values were used as the phenotype for association testing. Associations were evaluated using a linear mixed model to account for relatedness and population structure. | Protein levels were first rank-based inverse normal transformed, followed by regression to obtain residualized phenotypes. Residuals were adjusted for age, sex, sample collection site, and the first 10 genetic principal components. Genome-wide association analyses were conducted using BGENIE under an additive model, and results were combined via meta-analysis using METAL. | Normalized Protein expression levels were inverse-rank normalized. Discovery GWAS models included covariates for age, age<sup>2</sup>, sex, age-by-sex, age<sup>2</sup>-by-sex, batch, UK Biobank assessment centre, genotyping array, time between sampling and measurement, and the first 20 genetic principal components. Association testing was performed using REGENIE. |

We retained 516 proteins measured in the deCODE study that were matched to proteins in the brain pQTL dataset and showed genome-wide significant associations in both. Of these, 343 pairs (66.5%) demonstrated evidence of colocalization (posterior probability > 0.8; Figure 1A). Similarly, 529 proteins in the Fenland study and 540 proteins in UKB-PPP were matched to brain pQTLs, with 397 (75.1%; Figure 1B) and 435 (80.6%; Figure 1C) showing colocalization evidence, respectively. Detailed colocalization results are provided in Supplementary Table 1.

**Figure 1.**
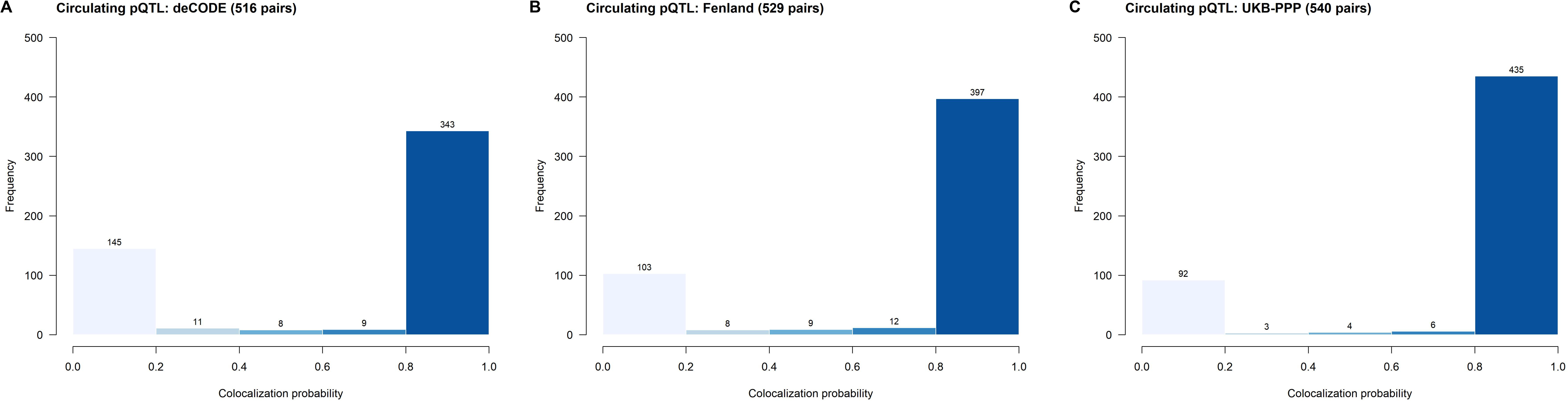
Distribution of colocalization probabilities between plasma and dorsolateral prefrontal cortex cis-pQTLs. Histograms show colocalization probabilities for circulating pQTLs derived from three cohorts: (A) deCODE, (B) Fenland, and (C) UKB-PPP. Counts above bars indicate the number of pairs per bin. A colocalization probability > 0.8 was considered evidence of colocalization.

Overall, genetic associations were largely consistent between plasma and brain among colocalized pQTLs, with Spearman’s rank correlation coefficients of 0.51 for deCODE, 0.53 for Fenland, and 0.47 for UKB-PPP. However, a notable proportion of colocalized pQTLs exhibited opposite effect directions between plasma and brain. Specifically, 67 pairs (19.5%; Figure 2A) in deCODE, 85 pairs (21.4%; Figure 2B) in Fenland, and 110 pairs (25.3%; Figure 2C) in UKB-PPP showed discordant effect directions (Supplementary Table 2).

**Figure 2.**
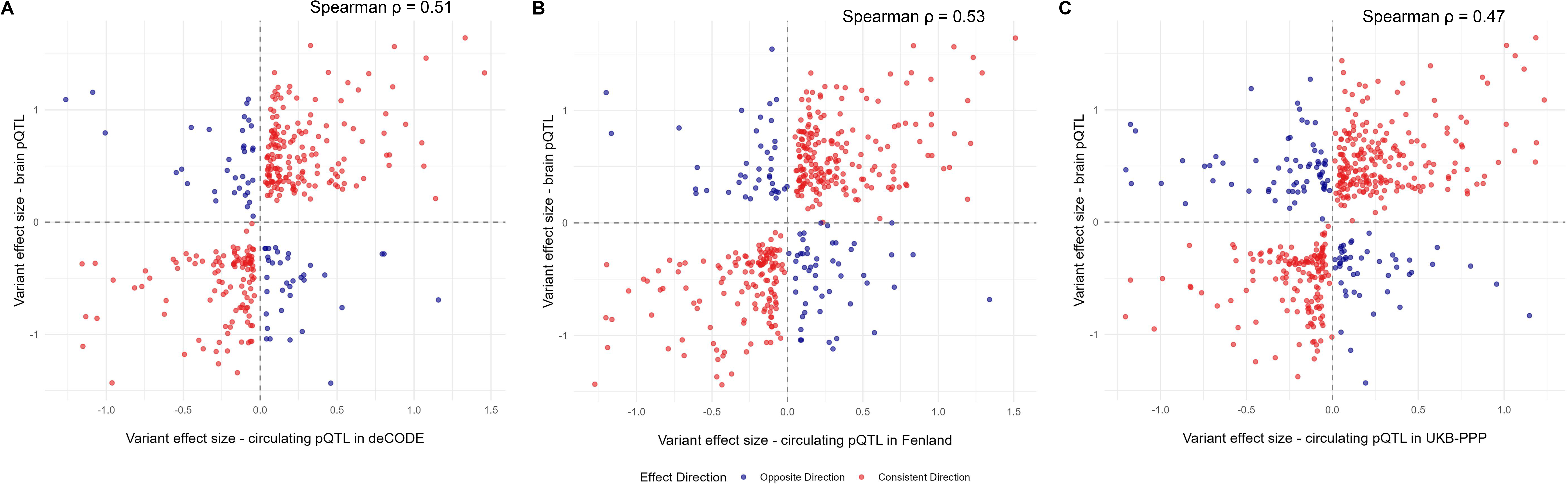
Comparison of variant effect sizes between circulating and brain pQTLs. Scatter plots illustrate the correlation of genetic effect sizes for pQTLs identified in dorsolateral prefrontal cortex and those in plasma across three cohorts: (A) deCODE, (B) Fenland, and (C) UKB-PPP. Red dots indicate concordant directions of effect across tissues, while blue dots indicate discordant directions. Dashed lines denote the null effect.

### Tissue-specific gene expression patterns

Tissue-specific gene expression enrichment z-scores are provided in Supplementary Table 3. Figure 3 depicts the proportion of genes with high tissue-specific expression stratified by colocalization status of circulating and brain pQTLs. Across all three circulating pQTL studies, proteins with non-colocalized pQTLs were more likely to exhibit higher gene expression levels in brain tissues compared to proteins with colocalized pQTLs. For instance, in the deCODE study, 31.21% of proteins with non-colocalized pQTLs showed high gene expression levels in the Brodmann area 9 (BA9), a key component of the dorsolateral prefrontal cortex, compared to 11.95% of proteins with colocalized pQTLs (Chi-squared test p-value = 1.90 × 10^-7^). Similar patterns were observed in the Brodmann area 24 (BA24; Chi-squared test p-value = 8.76 × 10^-6^), which is structurally and functionally linked to BA9, as well as across the broader cortex (Chi-squared test p-value = 5.36 × 10^-5^). These patterns were consistent based on the Fenland study and UKB-PPP (Figure 3).

**Figure 3.**
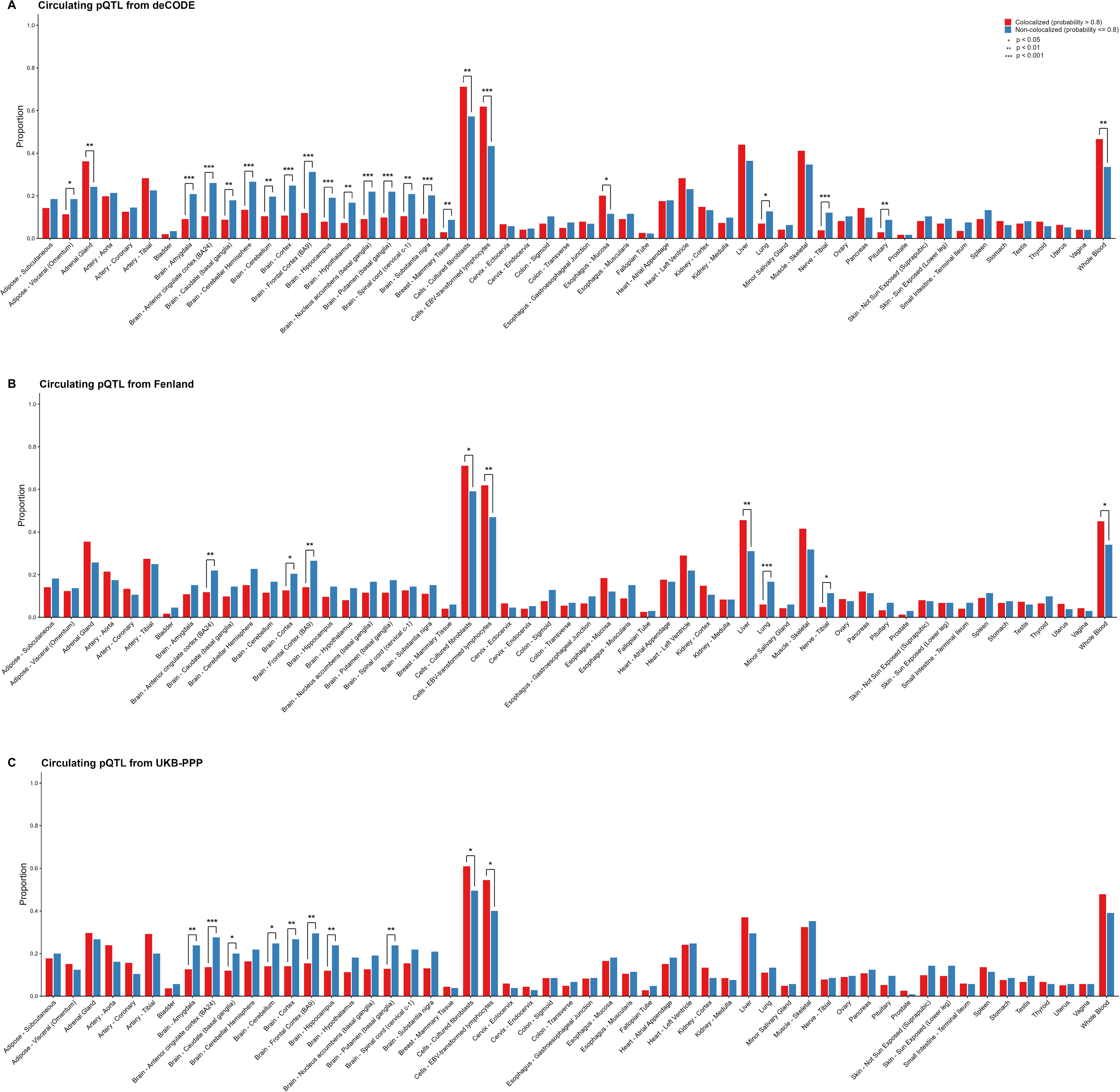
Tissue-specific gene expression patterns stratified by whether proteins have colocalized cis-pQTLs between plasma and dorsolateral prefrontal cortex. Genes with an enrichment z-score > 2 were defined as having elevated expression in a given tissue. Bars show the proportion of proteins with colocalized cis-pQTLs (red) or without colocalization (blue) that exhibit elevated expression in the corresponding tissue. Differences in proportions between groups were assessed using a Chi-squared test. Statistical significance is indicated as follows: *p < 0.05, **p < 0.01, and ***p < 0.001.

In contrast, proteins with colocalized pQTLs were more likely to have higher gene expression in non-neural tissues, including EBV-transformed lymphocytes, whole blood, cultured fibroblasts, and liver, which are involved in systemic protein production and immune function. For example, 61.81% of proteins with colocalized pQTLs showed high gene expression levels in EBV-transformed lymphocytes, compared to 43.35% of proteins with non-colocalized pQTLs (Chi-squared test p-value = 1.00 × 10^-4^).

### MR-estimated effects on neuroticism

Lastly, we performed MR analyses to evaluate whether discrepancies between plasma and brain pQTL effects could impact downstream analyses. Neuroticism was used as an illustrative outcome trait. The full MR test statistics are provided in Supplementary Table 4.

In total, 13 proteins showed significant associations with neuroticism (p-value < 1×10^-5^). Of these, seven proteins had consistent MR-estimated effect directions when instruments were derived from plasma and brain pQTLs: CD40, HSPA1A, ITIH3, MMAB, NCAM1, PMM1, and TMEM106B (Figure 4). For example, a one standard deviation increase in genetically predicted CD40 levels based on the UKB-PPP pQTL instrument was associated with a decrease of 0.037 (95% CI: 0.025–0.048) units in neuroticism score. A consistent association was observed when the dorsolateral prefrontal cortex pQTL instrument was used, where a one standard deviation increase in genetically predicted CD40 levels was associated with a decrease of 0.032 (95% CI: 0.022–0.042) units in neuroticism score.

**Figure 4.**
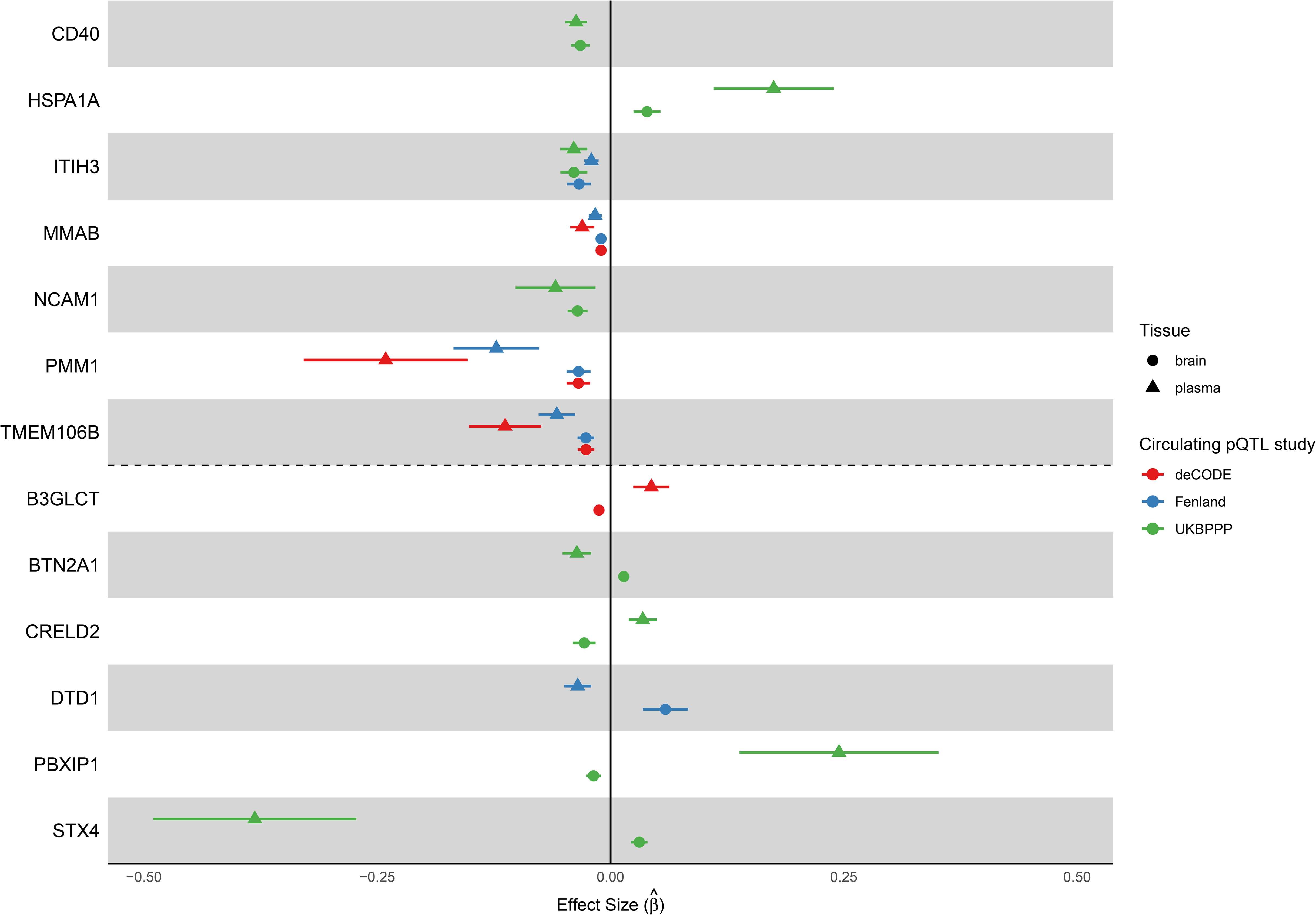
Significant associations between genetically predicted protein levels and neuroticism. Seven proteins showed consistent effect directions across instruments derived from different pQTL studies, while six showed discordant directions. Dots represent Mendelian randomization estimates, with error bars indicating 95% confidence intervals. Shapes denote the pQTL tissue source, and colors indicate the circulating pQTL discovery cohort. The vertical solid line indicates the null effect.

In contrast, six proteins showed discordant MR effect directions depending on whether plasma or brain pQTL instruments were used: B3GLCT, BTN2A1, CRELD2, DTD1, PBXIP1, and STX4 (Figure 4). For example, genetically predicted STX4 levels based on plasma pQTLs were associated with lower neuroticism scores (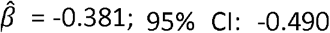 to -0.272), whereas the brain-derived instrument indicated an association with higher neuroticism scores (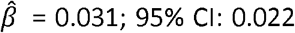 to 0.040). For B3GLCT, the plasma pQTL-based estimate suggested higher neuroticism scores (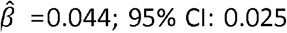 to 0.063), while the brain pQTL-based estimate suggested the opposite effect (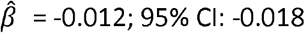 to -0.007).

## Discussion

Proteins play central roles in biological processes and disease mechanisms, and genetic studies of protein abundance have provided opportunities for identifying candidate biomarkers and therapeutic targets^36–38^. Large population studies have generated extensive pQTL resources based on circulating proteins, enabling integration of proteomic measurements with genetic association studies of complex traits^9–11^. However, circulating proteins may originate from multiple tissues and biological processes. As a result, genetic variants associated with circulating protein concentrations may not necessarily reflect the regulation of proteins within disease-relevant tissues. In this study, we combined a dorsolateral prefrontal cortex cis-pQTL dataset with three large circulating cis-pQTL resources to evaluate the consistency and discrepancies between genetic regulation of proteins in plasma and brain tissue by integrating colocalization analyses, tissue expression patterns, and MR analyses.

Across the proteins examined in both plasma and dorsolateral prefrontal cortex datasets, 66.5-80.6% of loci showed evidence of colocalization between circulating and brain pQTL signals, indicating that for many proteins, the same genetic variants influence abundance in both tissues. At the same time, a notable fraction of loci did not colocalize, reflecting tissue-specific genetic regulation for certain proteins. Among the colocalized loci, approximately 20 percent exhibited opposite directions of genetic effects between plasma and brain, suggesting that even when the same variant influences protein abundance, its effects on protein levels can differ across tissues.

Several biological mechanisms may contribute to these observations. Circulating protein measurements can reflect processes beyond steady-state intracellular abundance, including cellular turnover and tissue injury. For instance, variants that reduce protein abundance within cells could be associated with increased circulating levels if proteins are released into the bloodstream during cell death or degradation processes^39^. Conversely, variants that increase intracellular protein abundance may not lead to higher circulating concentrations when secretion is tightly regulated^40^. Such mechanisms may be particularly relevant for proteins expressed in the brain, where cellular damage or turnover could influence the presence of proteins in circulation^41,42^. These findings highlight the complexity of interpreting genetic associations with circulating protein levels and emphasize the importance of considering tissue context when evaluating pQTL signals. In addition, technical differences across studies, including variation in measurement platforms, detection sensitivity, and protein quantification methods, may also contribute to the observed discrepancies.

Our analysis of tissue-specific gene expression patterns further supports the presence of distinct biological origins for proteins detected in circulation. Proteins with colocalized pQTL signals between plasma and brain were less likely to show high gene expression levels in brain tissues and were more likely to be highly expressed in tissues involved in systemic protein production and immune function, including immune cells, fibroblasts, and liver. These tissues contribute substantially to the production of secreted proteins and circulating immune mediators, making it plausible that proteins originating from these sources would exhibit similar genetic regulation across plasma and brain samples. In contrast, proteins without colocalized pQTL signals were more likely to show high gene expression levels in brain tissues, including cortical regions closely related to the dorsolateral prefrontal cortex. This pattern suggests that proteins primarily produced within the brain may be subject to tissue-specific regulatory mechanisms that are not fully captured by circulating pQTL analyses.

To illustrate how these differences may influence downstream genetics-guided analyses, MR analyses were conducted using neuroticism as an illustrative outcome trait. Not all proteins demonstrating associations are expected to be brain-specific, as the brain proteome may also contain proteins originating from multiple sources, including peripheral immune cells and circulating proteins that cross the blood-brain barrier. Comparing MR estimates derived from circulating and brain pQTL instruments therefore provides a useful approach for evaluating the importance of tissue context.

Among the proteins showing significant MR associations with neuroticism, several exhibited consistent effect estimates regardless of whether circulating or brain pQTL instruments were used. One example is CD40, a receptor involved in immune activation and inflammatory signaling that is widely expressed in antigen-presenting cells and B cells^43^. Because CD40 functions primarily within immune pathways and circulating immune cells, it is plausible that circulating protein levels capture biologically relevant variation for this protein. TMEM106B also showed consistent estimated effects across tissues. TMEM106B encodes a lysosomal membrane protein^44^ that is highly expressed in neurons and oligodendrocytes^45^ and plays an important role in lysosome dynamics, including regulation of lysosomal pH, trafficking, and degradation pathways^44,46,47^, which are critical for neuronal homeostasis and have been implicated in brain aging and multiple neurodegenerative diseases such as frontotemporal lobar degeneration and related proteinopathies^48^. These functions suggest that genetic regulation captured by both circulating and brain pQTLs may tag biologically relevant processes affecting neuroticism⍰related pathways.

In contrast, several proteins showed discordant MR effect directions depending on whether circulating or brain pQTL instruments were used. For example, STX4 exhibited opposite estimated effects on neuroticism across tissues. STX4 encodes a member of the syntaxin family that participates in vesicle trafficking and membrane fusion processes that are important for neuronal signaling^49^. In this context, brain-derived genetic instruments may better capture biologically relevant regulation of this protein in neural tissues. Similar discrepancies were observed for B3GLCT, which encodes a glycosyltransferase involved in protein glycosylation pathways^50^. Although the specific functions of this protein in the brain remain incompletely characterized, the discordant estimates suggest that tissue-specific regulation may be important to consider when interpreting genetics-based analyses.

Not all discordant results indicate that circulating pQTLs lack relevance. For example, BTN2A1 showed opposite MR estimates between plasma and brain instruments, yet it is a key regulator of immune function. This gene, part of the butyrophilin family, interacts with V*γ*9V*δ*2 T⍰cell receptors to modulate *γδ* T⍰cell activation and cytokine responses, influencing systemic immune pathways^51^. Circulating protein levels may therefore capture biologically meaningful variation from immune processes even when brain-derived signals differ. This example highlights that the most informative data source may depend on the biological role and tissue origin of the protein under investigation, and that integrated evidence from multiple tissues and functional contexts can improve interpretation in genetics⍰guided analyses.

Several limitations of this study should be considered. First, our analyses were restricted to individuals of European ancestry because large-scale pQTL datasets in other populations remain limited. Second, our study focused on a single brain pQTL dataset due to the current scarcity of large-scale tissue-specific proteomic resources. Expanding pQTL analyses to additional brain regions and other tissues will provide a more comprehensive understanding of how genetic regulation of proteins varies across biological contexts. Third, the brain pQTL data were derived from post-mortem samples. Proteins measured in post-mortem tissues may be affected by processes such as protein degradation, enzymatic autolysis, and microbial activity that occur during the post-mortem interval^52,53^. In addition, we were unable to fully account for potential differences arising from technical variation across datasets, including differences in proteomic platforms, measurement protocols, and detection sensitivity. Finally, our analyses were limited to cis-pQTLs. We hypothesize that trans-pQTLs may exhibit greater discrepancies across tissues due to their stronger involvement in tissue-specific regulatory mechanisms. Continued efforts to characterize proteomic variation across tissues, particularly in non-blood samples, will be important for improving the interpretation of genetic associations with protein abundance.

In summary, our findings demonstrate that circulating pQTL signals do not always reflect the genetic regulation of proteins within disease-relevant tissues. Even when genetic associations colocalize between plasma and brain, the direction of genetic effects may differ, which can influence downstream analyses. At the same time, circulating pQTL resources remain valuable for studying proteins originating from systemic tissues and immune pathways. As larger proteomic datasets across multiple tissues become available, integrating tissue-specific genetic regulation with circulating protein measurements will improve our ability to interpret genetic associations and identify biologically meaningful biomarkers and therapeutic targets.

## Statements & Declarations

### Author Contributions

T.L. conceptualized the study. T.L. and W.Z. designed the methodology. Y.C. and T.L. acquired the data. Y.C. performed data analysis. Y.C. wrote the initial draft of the manuscript. All authors interpreted the results, revised the manuscript critically, and approved the final version of the manuscript.

### Data Availability

The pQTL summary statistics from the deCODE study are available at https://www.decode.com/summarydata/. The dorsal prefrontal cortex pQTL summary statistics (syn64600176) and the pQTL summary statistics from the Fenland study (syn51761394) and the UKB-PPP (syn51364934) are available at https://www.synapse.org/ under respective accession numbers. All other data supporting the findings of this study are available within the paper and its Supplementary Information.

### Ethics approval

This study did not involve human or animal subjects.

### Consent to participate

Not applicable.

### Consent to publish

Not applicable.

